# Structure of a Ca^2+^ bound phosphoenzyme intermediate in the inward-to-outward transition of Ca^2+^-ATPase 1 from *Listeria monocytogenes*

**DOI:** 10.1101/2024.03.06.583647

**Authors:** Sara Basse Hansen, Rasmus Kock Flygaard, Magnus Kjaergaard, Poul Nissen

## Abstract

Active transport by Ca^2+^-ATPases of the P-type ATPase family maintain a very low cytosolic calcium concentration and steep electrochemical gradients. Detailed mechanisms of this transport have been described from structures of mammalian sarco/endoplasmic reticulum Ca^2+^-ATPases (SERCA) stabilized by inhibitors at specific intermediate steps of the transport cycle. An essentially irreversible step is crucial to prevent reflux in active transport against steep gradients. Single-molecule FRET (smFRET) study of the bacterial Ca^2+^-ATPase LMCA1 revealed an intermediate of the transition between so-called [Ca]E1P and E2P states, suggesting that calcium release from this intermediate is the irreversible step. Here, we present a 3.5Å cryo-EM structure for a four-glycine insertion mutant (G_4_-LMCA1) in a lipid nanodisc obtained under turnover conditions and adopting such a calcium-bound intermediate, denoted [Ca]E2P. The cytosolic domains are positioned in the E2P-like conformation, while the calcium-binding transmembrane (TM) domain is similar to calcium-bound E1P-ADP like conformation of SERCA. Missing density for the E292 residue at the calcium site (equivalent of SERCA1a E309) suggests flexibility and a site poised for calcium release and proton uptake. The structure suggests a mechanism for the inward-to-outward transition in Ca^2+^-ATPases, where ADP release and re-organisation of the cytoplasmic domains precede calcium release.

## Introduction

Cell viability depends on the interplay between many different pathways that are regulated and interconnected by e.g. metabolic or cell cycle feedback and environmental conditions. Changes in calcium concentration form potent signals in nearly all cells, and it is crucial to understand how calcium homeostasis is maintained and regulated. Ca^2+^-ATPases transport calcium out of the cell or into internal stores and maintain a low cytoplasmic calcium concentration that can be four orders of magnitude below surroundings^1,2^. Considering also a typically negative membrane potential and the divalent charge of calcium ions, cell membrane calcium gradients are some of the most powerful electrochemical gradients known in biological systems. As a consequence, Ca^2+^-ATPase transport against such gradients must include a practically irreversible step to avoid reflux. Structural and mechanistic examination of this irreversible step has remained elusive, but could explain key principles of active transporters.

The P-type ATPase family encompasses integral membrane proteins that share both structural and mechanistic features associated with large domain re-arrangements during the transport cycle^3^. P-type ATPases perform their active transport with energy from ATP hydrolysis via autoformation and breakdown of a phosphoenzyme^4^ and with structural transitions alternating between inward- and outward-oriented states^5,6^. Ca^2+^-ATPases encompass the P2A and P2B subtypes of the P-type family^7^, consisting of typically ten membrane-spanning helices, M1-10, connected to three cytosolic domains: Nucleotide (N) binding, phosphorylation (P) and actuator (A) domain, where the latter is involved in dephosphorylation and contains an essential TGES loop^8^. The mammalian sarco/endoplasmic reticulum membrane (SERCA) is highly studied and often serves as a general model^9^. SERCA explores inward-open (E1) and an outward-open (E2P) conformations, exposing the ion binding sites in the middle of the TM domain to either side of the membrane^6,10,11^. In the E1 state, two calcium ions enter the binding site via an entry pathway formed by M1-2, M4 and M6. Following Ca^2+^ binding, reorientation of the ATP-bound N domain facilitates transfer of the γ-phosphate from ATP to a catalytic aspartate in the P domain. Along with phosphorylation forming the E1P state, the ion pathway closes and bound calcium becomes occluded^12,13^. Next, the A domain makes a large rotational movement perpendicular to the membrane, and the N domain moves away from the P domain in formation of the outward-open E2P state. Altogether, these domain movements trigger opening of the calcium exit pathway formed by M1-6. Calcium is released to the other side of the membrane and protons occupy the ion binding sites in the E2 state^14^, where a conserved TGES loop from the A domain is now positioned to catalyze dephosphorylation of the aspartyl phosphorylation site. Dephosphorylation allows an E2-E1 transition with the E1 state releasing counter-transported proton to the cytoplasmic side and opening the cytoplasmic entry pathway^15^. However, key steps and intermediates are not yet accounted for with structural information in the [Ca_2_]E1P-ADP to E2P transition. When are ADP and calcium ions released, new ATP bound – and in which order, and how is an essentially irreversible step introduced?

In SERCA, a four-glycine insert in the A-M1 linker (G_4_-SERCA) allows occlusion of calcium in a stalled [Ca_2_]E2P intermediate state^16,17^. A similar intermediate was characterized by single-molecule Förster Resonance Energy Transfer (smFRET) in the equivalent G_4_ construct of the bacterial homolog Ca^2+^-ATPase 1 from *Listeria monocytogenes* (G_4_-LMCA1), where smFRET traces indicated a calcium-bound, but ADP-insensitive [Ca]E2P state with a partially rotated A domain^18^. Subsequently, the A domain makes the final rotation in formation of the outward-open E2P, which once reached showed no signs of reversal to the [Ca]E2P state. Hence, calcium release from the [Ca]E2P intermediate state was described as the critical, irreversible step. It points also to the A-M1 linker as crucial for transferring the energy from ATP hydrolysis to extracellular release of calcium, as was also observed for SERCA^16,17^. Other recent studies have described intermediate states for SERCA through time-resolved x-ray scattering (TR-XSS)^19^ and molecular dynamics (MD)^20^, generally confirming the rotation of the A domain as an important reaction coordinate for the E1P-E2P transition.

LMCA1 shares 34-39% sequences identity with mammalian SERCA (ATP2A genes), but it transports only a single calcium ion^21^ for each transport cycle (Figure 1A), similar to the plasma membrane Ca^2+^-ATPase (PMCA, ATP2B genes)^22^ and the secretory pathway Ca^2+^-ATPase (SPCA, ATP2C genes)^23,24^ that are also closely related although with a slightly lower sequence identity to LMCA1 (30-34%). We have previously reported crystal structures of LMCA1 stalled in presumably proton-occluded [H]E2-BeF_x_ and [H]E2-AlF_x_ forms and also G_4_-LMCA1 in the [H]E2-BeF_x_ form^25^. These structures were associated with a closed extracellular pathway and a TGES motif positioned for dephosphorylation (hence presumed all to be proton-occluded forms), which is unlike the SERCA E2-BeF_x_ form with an outward-open calcium exit pathway. The structures were consistent with sequence alignments^25^ to indicate a single calcium binding site (protonated, not calcium bound) equivalent of the calcium binding site II of SERCA, whereas the region corresponding to SERCA site I is occupied by Arg795 in LMCA1.

**Figure 1.**
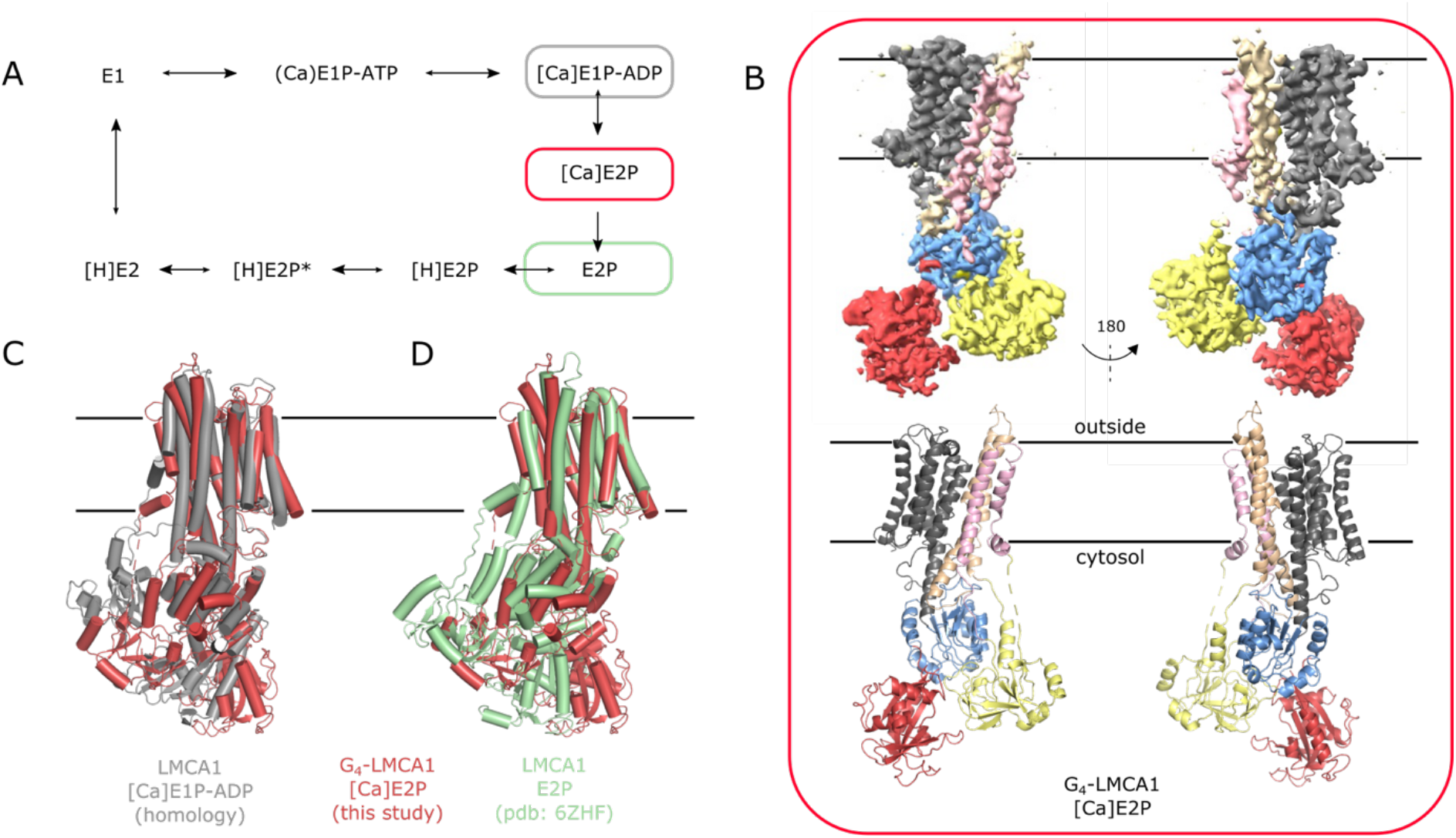
The cryo-EM structure of G_4_-LMCA1 adopts an intermediate E1P to E2P conformation. **(A)** Functional transport cycle of LMCA1. **(B)** 3.5Å cryo-EM map and model of G_4_-LMCA1 in the [Ca]E2P state. The map and model are color coded according to the domains (N red, A yellow, P blue, M1-2 lightpink, M3-4 wheat, M5-10 grey). **(C)** Alignment of [Ca]E2P (colored according to (B)) with an [Ca]E1P-ADP LMCA1 homology model based on SERCA [Ca_2_]E1-AlFx-ADP crystal struture (pdb: 1t5t) (grey). **(D)** Alignment of [Ca]E2P (colored according to (B)) with LMCA1 E2-BeFx crystal structure (pdb: 6zhf) (green). The structures are aligned at TM7-10. The volume made in ChimeraX^40^ and models are made in PyMOL.

Here, we present a cryo-EM structure at 3.5 Å resolution for G_4_-LMCA1 obtained under ATP turnover conditions and showing bound calcium at the predicted site. The intermediate represents a new structure for Ca^2+^-ATPases and ion-transporting P2-type ATPases in general, and is overall a hybrid between E1P and E2P states and a presumed intermediate in the E1P-E2P transition preceding the irreversible state of ion release.

## Results

LMCA1 was reconstituted in salipro nanodiscs^26^, which serves both to embed the protein in a lipid environment and remove the detergent background signal in EM. Wild-type (WT) LMCA1 remains active when reconstituted in salipro nanodiscs as assessed by a phosphate-release assay (Figure S1), albeit with a slower rate compared to the detergent-solubilized form. In detergent, G_4_-LMCA1 shows lower activity compared to WT^18^, but surprisingly, no P_i_ release activity could be measured for saposin-solubilized G_4_-LMCA1 (Figure S1). This may indicate that the saposin nanodisc stabilizes a particular state occurring in the ATPase reaction for LMCA1 and in particular associated with the G_4_ construct. Knowing these preconditions, we proceeded with structural analysis of the sample.

### Overall conformation of cryo-EM structure

The published smFRET data for detergent-solubilized G_4_-LMCA1 supplemented with 1 mM Ca^2+^ and 1 mM ATP revealed a predominant cycling between an E1-ATP and [Ca]E2P state showing high and medium-low FRET efficiency, respectively^18^. To catch the [Ca]E2P state for cryo-EM studies, we incubated saposin-solublized G_4_-LMCA1 with 1 mM Ca^2+^ and 1 mM ATP. The sample was incubated at room temperature for 2-10 minutes before plunge-freezing and cryo-EM imaging. Expecting two or more conformations from the cryo-EM dataset, both 3D classification and 3D variability analyses were applied in a search for several conformations of G_4_-LMCA1. However, only a single conformation was found after thorough processing of the data (Figure 1B) and it showed clear indications of calcium binding and a hybrid conformation between E1P and E2P conformations. So far, no structures have been reported for LMCA1 in E1 states, so we constructed a homology model based on the [Ca_2_]E1-AlF_x_-ADP structure of SERCA (based on pdb entry 1t5t)^12^. Previously, we published the crystal structure of the E2P state of LMCA1 (based on an E2-BeF_x_ form, pdb: 6zhf)^25^.

The structure of G_4_-LMCA1 obtained here represents a new conformation, from now on termed [Ca]E2P, that overall is distinct from both the [Ca_2_]E1-AlF_x_-ADP form of SERCA1a and a calcium-free E2P state of LMCA1 (RMSD values overall of 7.6 Å and 4.2 Å, respectively, Table S1). Still, a RMSD of 2.9 Å for the cytoplasmic domains comparing the G_4_-LMCA1 [Ca]E2P and the E2-BeF_x_ form of LMCA1 indicates a close similarity of the cytoplasmic headpiece. Similarly, an RMSD values of 2.1 Å between the TM domain of the G_4_-LMCA1 [Ca]E2P structure and SERCA1a [Ca_2_]E1-AlF_x_-ADP ^12^ indicates a close similarity of the calcium-bound TM-domain. Further visual inspection reveals closed ion pathways, and density consistent with a single calcium ion bound and occluded at a binding site between M4 and M6 (Figure 2B). Hence, the TM domain adopts an calcium-occluded E1P-like conformation, but the position of the M1-4 bundle is shifted relative to the [Ca_2_]E1-AlF_x_-ADP state of SERCA1a (Figure 1C). Indeed, the cytosolic domains, although configured in an E2P-like conformation, are not yet tilted relative to the membrane plane, which is otherwise typical of the E2P state (Figure 1D). Phosphorylation of the catalytic aspartate (Asp334) in the P domain is clearly visible (Figure S3), and the phosphorylation site is rotated away from the nucleotide binding site of the N domain. Visual inspection of the cryo-EM density map indicates no density for bound nucleotide at the N domain despite a 1 mM background of ATP/ADP in the cryo-EM sample. Altogether, these observations point to an intermediate state of the transition between the inward-occluded [Ca]E1P-ADP state to the outward-open E2P state, where ADP has been released, but calcium is still bound. Although the conformation displays both E1P and E2P features, we denote it a [Ca]E2P state to indicate the calcium-bound conformation of the transmembrane domain and the E2P-like, ADP insensitive conformation of the cytoplasmic domains.

**Figure 2.**
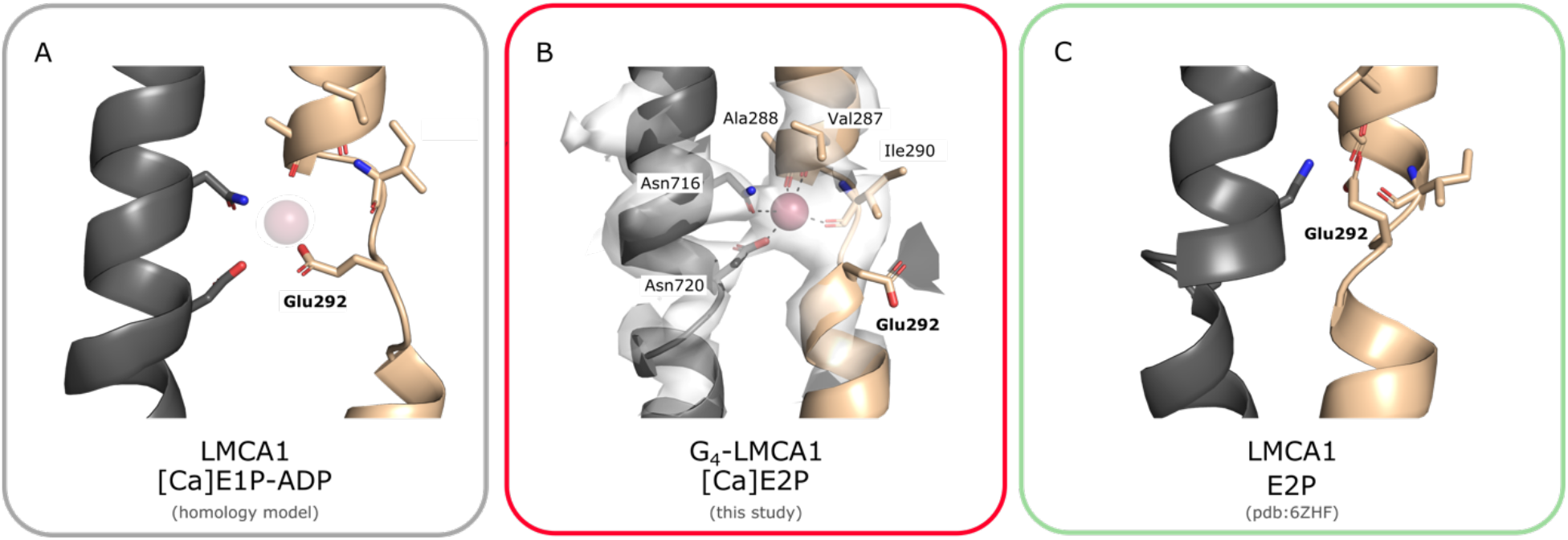
Glu292 coordination varies in different states. The calcium binding site between M4 and M6 is shown for **(A)** the [Ca]E1P-ADP LMCA1 homology model based on SERCA [Ca_2_]E1-AlF_x_-ADP crystal structure (pdb: 1t5t), (B) the [Ca]E2P structure, and **(C)** the LMCA1 E2-BeF_x_ crystal structure (pdb: 6zfh). In **(B)** the electron density map is shown for M4, M6, and the calcium ion at contour level 10. Calcium coordinating residues are shown as sticks.

### The calcium ion is bound at a flexible site

The structure of G_4_-LMCA1 in the [Ca]E2P state is consistent with the suggested calcium binding site with a bound calcium ion between M4 and M6, equivalent of SERCA site II (Figure 2B). The calcium ion appears to coordinate two side chains in M6 (Asn716 and Asn720) and backbone carbonyl groups in M4 (Val287, Ala288, Ile290). Water molecules are likely also part of the coordination, although they could not be resolved in the 3.5 Å resolution cryo-EM map. For the [Ca]E1P-ADP homology model of LMCA1, Glu292 would point towards the calcium ion, as for SERCA1a (Figure 2A). However, the map for the [Ca]E2P form of G_4_-LMCA1 reveals no density for Glu292 oriented towards the calcium ion (Figure 2B). Rather, Glu292 appears flexible and pointing away from the site. In the [H]E2P-like structure of LMCA1, representing a subsequent calcium-released form, Glu292 occupies the vacated calcium binding site, and prediction of a high pK_a_ indicates that it is most likely protonated^25^ (Figure 2C).

### Conformational changes for the [Ca]E1P-ADP to E2P transition

ATP hydrolysis overall drives the conformational changes that lead to calcium translocation, and we will focus here on the transitions following the phosphorylation step, in particular the [Ca]E2P to E2P transition of LMCA1, which from smFRET data appears as the critical, irreversible step of calcium transport.

In the [Ca]E2P form of G_4_-LMCA1, the P domain is placed at an intermediate position relative to the TM domain (Figure 3E), and the A and N domain point away from the membrane, which puts G_4_-LMCA1 in an upright conformation as compared to [Ca]E1P-ADP (Figure 3B) and [H]E2P (Figure 3H). The A domain is partially rotated towards the position of the E2P state (Figure 3A,D,G). The nucleotide binding site is empty; ADP must have been released prior to this point, probably at a preceding step going from the [Ca]E1P-ADP state to an ADP-sensitive [Ca]E1P state^27^.

**Figure 3.**
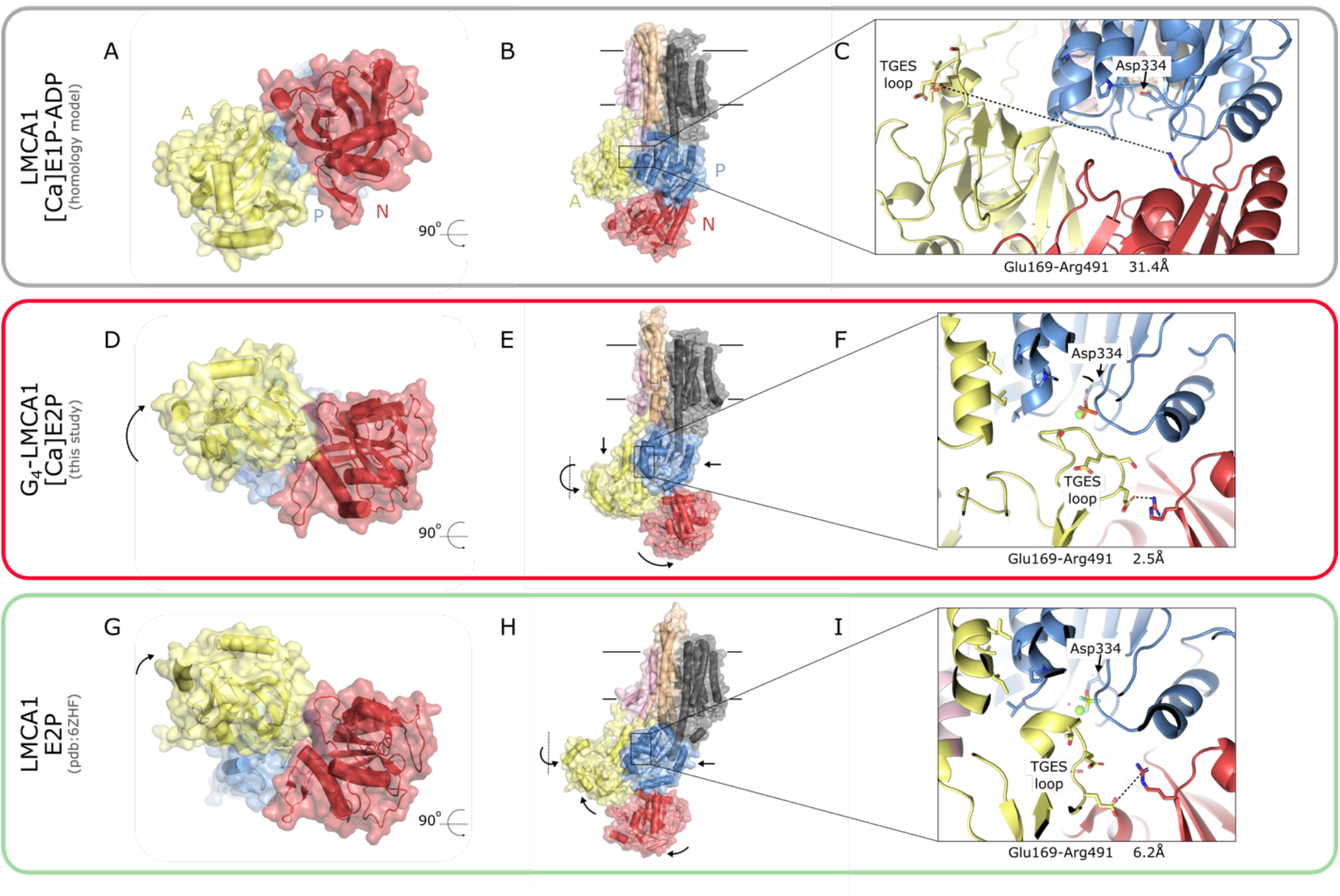
Principal movements of cytosolic domains LMCA1 during the E1P to E2P transition. Surface and cartoon representation of LMCA1 (**A**,**B**,**C**) [Ca]E1P-ADP LMCA1 homology model based on SERCA [Ca_2_]E1-AlF_x_-ADP crystal structure (pdb: 1t5t), (**D**,**E**,**F**) the G_4_-LMCA1 [Ca]E2P structure, and (G,H,I) the LMCA1 E2-BeF_x_ crystal structure (pdb: 6zfh). The structures are colored according the Figure 1. Arrows indicate forward domain movements. (**A**,**D**,**G**) The structures represent the rotation of the A domain relative to the P domain. (**B**,**E**,**H**) The structures represent the movement of the cytosolic domains. Zoom-in panels show the TGES loop away from the phorphorylation site and a large distance between Glu169 and Arg491 (**C**), the TGES loop protecting the phosphorylation site and Glu169-Arg491 interaction (**F**), and the TGES loop primed for dephosphorylation and a break in the Glu169-Arg491 interaction (**I**). Hydrophobic interacting residues (Ile218, Leu221, Leu222 in the A domain and Pro601, Val625, Pro629 in the P domain) between the A and P domain are shown as sticks in (F,I).

In the [Ca]E2P intermediate of G_4_-LMCA1, the A domain is rotated to allow interaction with the P domain through hydrophobic interactions between the A and P domain (Figure 3F,I). A salt bridge between the A domain (Glu169) and the N domain (Arg491) stabilizes the A domain in its intermediate position (Figure 3F). When the A domain rotates further in the E2P state, this interdomain bond is disrupted, but the hydrophobic interactions between the A and P domain remain (Figure 3I). In the [Ca]E2P state, the TGES loop in the A domain blocks access to the phosphorylated aspartate in the P domain for a reverse reaction with ADP (Figure 3F). However, unlike the calcium released E2P state, where the pump is primed for dephosphorylation^25^, the TGES motif of [Ca]E2P is not yet positioned for catalysis at the phosphorylated aspartate (Figure 3I).

### The A-M linkers couple cytosolic domain to TM movement

Energy deposited by ATP hydrolysis must be transmitted during the [Ca]E1P-ADP to E2P transition. Both biochemical and computational studies suggest that energy is stored in the linkers between the A domain and the transmembrane helices M1 and M2 (and M3) and that the strain built up transmits to the opening of the TM domain and release of calcium^17,20^.

Figure 4 highlights the variation of length spanned and conformation of the A-M1 and M2-A linkers. The A-M1 linker is unstructured without any apparent interactions in [Ca]E1P-ADP (model based on SERCa1a crystal structures) and in the crystal structure of E2-BeF_x_ representing [H]E2P state of LMCA1. The same is true for the [Ca]E2P intermediate state with an A-M1 linker that is extended by four glycines. However, the distance spanned by this linkers is almost 6Å longer than in the [Ca]E1P-ADP and E2P states (Figure 4A,B,D) and without the G_4_ insert it would not reach without undergoing some level of conformational changes (Figure S4). Furthermore, four hydrophobic residues (Leu104, Ala106, Leu107, Met110) in a helical part of the M2-A linker interact in a hydrophobic network with M4 and the P domain in the modelled [Ca]E1P-ADP state of LMCA1 (based on SERCA1a [Ca_2_]E1-AlF_x_-ADP, Figure 4A). This network is disrupted in the [Ca]E2P state, where the helix is unwound and stretched by almost 10Å. This change allows the A domain to obtain a partially rotated position in the [Ca]E2P state (Figure 4B). The four hydrophobic residues in the M2-A linker twist away from the P domain interface, and instead an ionic interaction network is formed between M2-A and the P domain (Figure 4C) stabilizing the M2-A linker in an extended conformation. In the calcium released E2P state, M1-2 are rotated and the M2-A linker again shortened by helix formation, incorporating the four hydrophobic residues that in the [Ca]E2P intermediate form a hydrophobic network with the A domain (Figure 4D).

**Figure 4.**
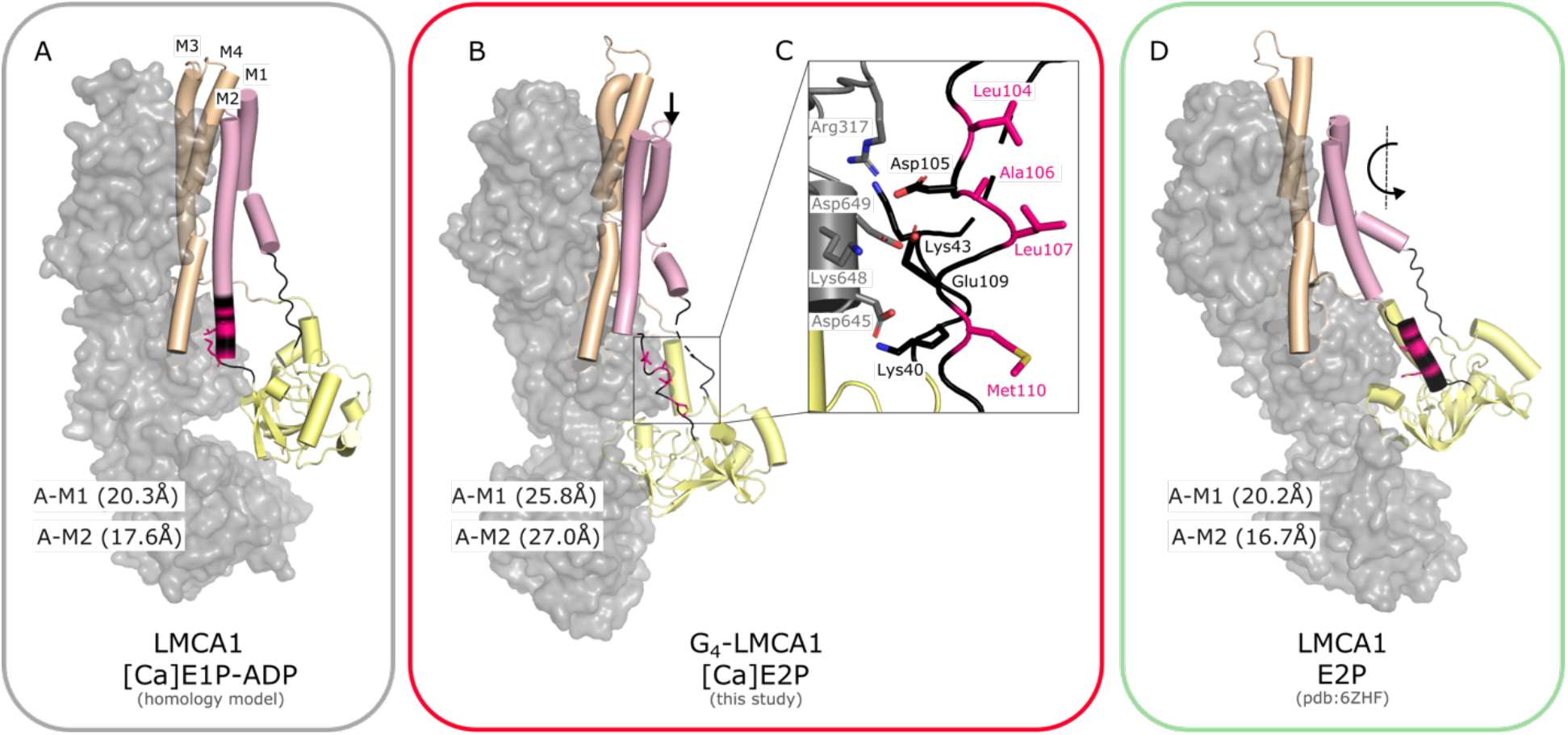
A-M2 linker changes its conformation during the E1P to E2P transition. Surface and cartoon representation of LMCA1 **(A)** [Ca]E1P-ADP LMCA1 homology model based on SERCA [Ca_2_]E1-AlF_x_-ADP crystal structure (pdb: 1t5t), **(B)** the G_4_-LMCA1 [Ca]E2P structure, and **(D)** the LMCA1 E2-BeF_x_ crystal structure (pdb: 6zfh). The A domain and M1-4 are shown as cartoons and colored acoording to Figure 1. M5-10, the N and P domain are shown as a grey surface. The A-M1 linker (Leu39-Pro46) and the M2-A linker (Ser102-Pro113) are colored black, and the distance between C_α_Leu38-C_α_Pro46 and C_α_Ser102-C_α_Pro113, respectively, is shown. The four hydrophobic residues (Leu104, Ala106, Leu107, and Met110) are clored pink. Arrows indicate forward domain movements. **(C)** Zoom-in panel shows ionic interactions between A-M2 and residues in the P domain.

## Discussion

During the transition from calcium-occluded E1P-ADP to calcium released E2P of Ca^2+^-ATPases, several things happen: ADP is released, the cytoplasmic domains rotate, and altogether they tilt relative to the membrane, the TM domain opens, and calcium is released. We have determined a structure of the elusive calcium-occluded, ADP insensitive E2P intermediate, [Ca]E2P, for LMCA1 by cryo-EM using a G_4_ insertion mutant form known to pause at this intermediate. It sheds light on the sequence and mechanism of structural changes associated with calcium release. The [Ca]E2P structure has a TM domain that adopts an E1P-like conformation with bound calcium; for LMCA1 a single calcium ion at the site corresponding to site II in SERCA. The cytosolic domains adopt an E2P-like configuration with a partially rotated A domain, and the ADP nucleotide is released. However, the cytoplasmic headpiece of the three cytoplasmic domains is still not tilted into the configuration of the E2P state, but instead paused in the intermediate state due to the extension of the A-M1 linker with the G_4_-insert that relaxes strain of the A-M1 linker that otherwise would transmit energy into formation of a calcium-released E2P state

### A transient interaction in the E1-E2 transition

Several transient conformations connect ATP hydrolysis to calcium release, which is illustrated for LMCA1 in Figure 5. Upon phosphoryl transfer from ATP and ADP release, the cytosolic domains re-orientate into an E2P-like configuration leading to the [Ca]E2P intermediate. Here, the TGES loop in the A domain shields the phosphorylation site in the P-domain and the empty nucleotide binding site in the N-domain, and the A-N interdomain interaction are stabilized by an ionic interation between the A and N domain, Glu169 and Arg491 respectively (Figure 3F). Arg491 in LMCA1 is conserved in all P2-type ATPases and corresponds to Arg560 in SERCA, which is indeed important for nucleotide binding^12,13,28^. An Arg560Ala mutation is known to stimulate the transition to the calcium-free E2P state^29^, which could be explained by loss of the interaction that stabilizes the preceding state. This suggests that Arg491 in the N domain has a dual role of nucleotide binding in the [Ca]E1-ATP and [Ca]E1P-ADP states, and thereafter A domain stabilization in the transient [Ca]E2P state that prepares for Ca^2+^ release. In the [Ca]E2P state, the calcium site features a flexible Glu292 residue (as indicated by poor density and an orientation away from the Ca^2+^ site) and hence lower coordination by protein side chains, which may prime solvation and release of the Ca^2+^ ion in the subsequent step. At the same time, the Glu292 side chain can pick up a proton, presumably from the extracellular environment, and enter the vacant site in a neutral form after Ca^2+^ release, i.e. leading to proton occlusion and dephosphorylation. Glu292 corresponds to Glu309 in SERCA, which again is conserved in all P2-type ATPases; essential in calcium binding and gating^30-32^. Contrary to wildtype ATPases, the [Ca]E2P state of G_4_-LMCA1 is relaxed by the four-glycine insert in the A-M1 linker, slowing down a final transition to release calcium, and therefore it resides in a [Ca]E2P state as revealed here by cryo-EM under ATPases conditions in a lipid nanodisc. The G_4_-LMCA1 structure shows that the A-M1 linker must undergo strain in the wild-type [Ca]E2P form. The linker plays a crucial role in formation of a calcium exit pathway, which is in agreement with already published experiments on phosphorylation and calcium release kinetics^16,17^, smFRET^18^ and molecular dynamics^20^. Figure S4 shows density of the A-M linkers of G_4_-LMCA1 [Ca]E2P. Parts of the M2-A linker are clearly defined in density and dock to the P domain, and it indicates that these interactions too stabilize the [Ca]E2P state. Presumably, both the A-M1 and the M2-A linker stretch in the wild-type [Ca]E2P intermediate and “pull open” a calcium exit pathway. For SERCA this open state forms a stable intermediate, whereas for LMCA1 it will immediately close with proton occlusion, most likely associated with a very high pK_a_. Unlike for SERCA, calcium release in LMCA1 therefore is associated with direct rotation of the A domain in formation of a closed E2P state, [H]E2P, likely with occlusion of a proton picked up at Glu292 in a direct pathway, as also indicated earlier by crystal structures^25^. In contrast, SERCA likely features a back-door pathway for proton entry^33^ and explores an outward-open E2P prior to proton occlusion at two co-operative sites.

**Figure 5.**
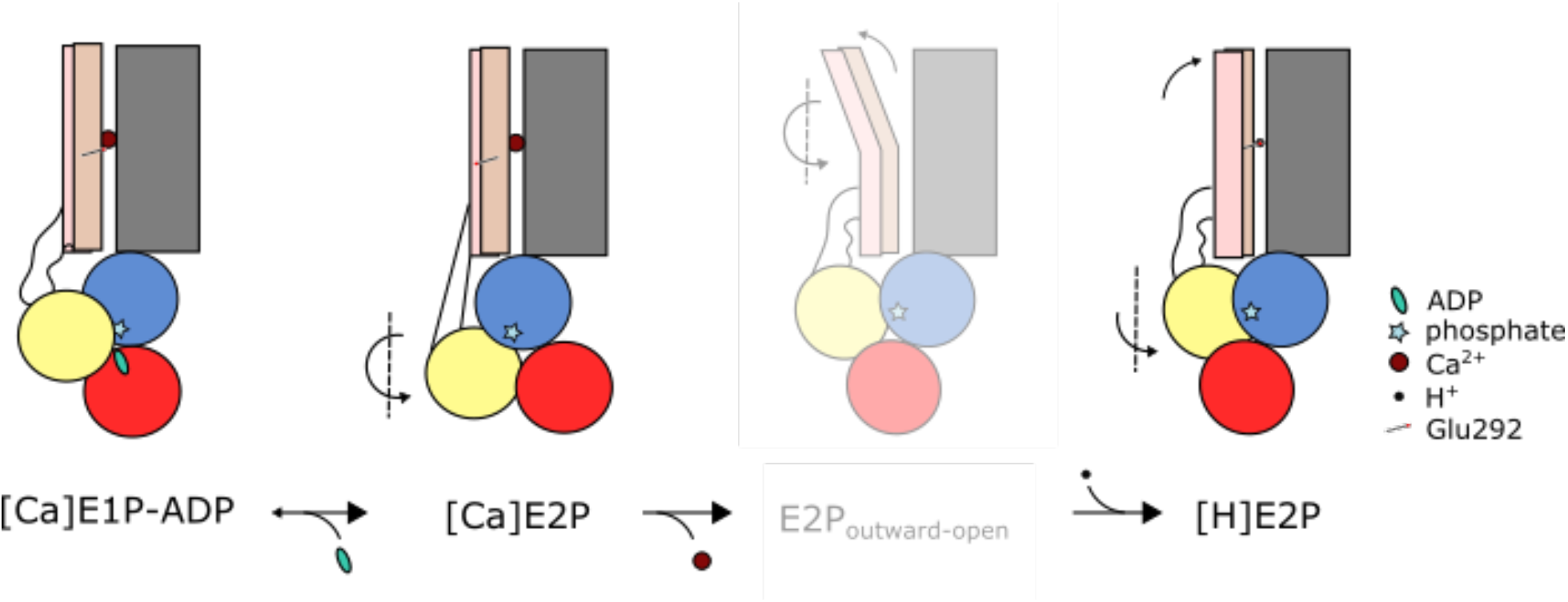
Sequence of events in the E1-E2 transition. Schematic representation of the intermediate conformations during the E1-E2 transition. The E2P_outward-open_ conformation is blurred to indicate that it is transient for LMCA1.

### Calcium release is the irreversible step for LMCA1

To transport calcium ions against steep concentration gradients, at least one step in the reaction cycle must be practically irreversible and therefore resistant to the gradient. Earlier smFRET studies^18^ and the studies presented here indicate that the LMCA1 [Ca]E2P state is the critical intermediate. The A domain completes its rotation only when calcium is released and this transition appears irreversible^18^, presumably linked to very fast kinetics of occlusion, as decribed above, and the subsequent dephosphorylation reaction of LMCA1 once the E2P state is reached. Indeed, our crystal structure of the E2-BeF_x_ state shows that the enzyme is primed for dephosphorylation with the glutamate side chain of the TGES already positioned over the phosphorylation site to catalyze the in-line attack of a water molecule^25^. The G_4_-LMCA1 E2-BeF_x_ form does not enter a [Ca]E2P like state in reverse reaction with Ca^2+^ present^18,25^, unlike what SERCA has been reported to do^16,17^. The outward-open E2P state therefore is short-lived for LMCA1, and the kinetics are too fast to determine a single irreversible step. Our analysis of the G_4_-LMCA1 [Ca]E2P state confirms that a final A domain rotation must happen in transition to the [H]E2P state, but also that the opening/closing of the calcium exit pathway occur during this transition. Hence the irreversible transition for LMCA1 is [Ca]E2P to [H]E2P, which completes the the A domain rotation transmitted by linker regions, and which includes calcium release with no outward-open Ca^2+^ intermediate explored *per se*.

The minimal exposure of an outward-open state is a powerful mechanism that avoids reversible function in steep electrochemical gradients for Ca^2+^, and it may also protect the cell against inhibitors of the Ca^2+^-ATPase from the surrounding environment that may find a binding site in the outward-open pathway. The LMCA1 studies presented here add valuable information on both P-type ATPase mechanisms in general and Ca^2+^-ATPases in bacteria specifically.

## Materials and methods

G_4_-LMCA1 was constructed, expressed, and purified using His-tag affinity and size-exclusion chromatography as described earlier^25^.

### Reconstitution in salipro nanodiscs

75 μl of detergent-solubilized G_4_-LMCA1 at 10 mg/ml in buffer (50 mM Tris, 200 mM KCl, 10 mM MgCl_2_, 20% v/v glycerol, 1 mM BME, 0.25 mg/mL C_12_E_8_, pH=7.6) was mixed with 75 μl brain extract lipids at 5 mg/ml and incubated at room temperature for 30 min. The brain extract lipids were prepared at 20 mg/ml in buffer (1.5% w/v DDM, 50 mM Tris-HCl, 150 mM KCl, pH 7.5) and diluted four-fold to obtain a concentration of 5 mg/ml. 250 μl of saposin A at 5 mg/ml in buffer (20 mM HEPES, 150 mM NaCl, pH 7.5) was added to the mix and incubated for 10 min at room temperature. Saposin A was expressed and purified according to earlier reports^34^. The reconstituted monomeric fraction was separated from aggregates and empty discs on a Superdex 200 10/300 in buffer (50 mM Tris-HCl, 150 mM KCl, pH=7.6).

Reconstitution of WT LMCA1 used for ATPase activity assay was carried out similar to G_4_-LMCA1, but worked best in small reaction volumes and 17 parallel reactions with 5 μl protein, 5 ul brain extract lipids and 20 μl saposin A was mixed under the same conditions. After reconstitution, the small reactions were pooled, and the monomeric fractions were collected from a Superdex 200 10/300. The results are shown in Figure S1.

### ATPase activity assay

The ATPase activity was measured by the Baginski method^35^ with quantification of inorganic phosphate released by LMCA1 after reaction with CaCl_2_ and ATP. The reaction mix was started by addition of ATP (25 mM stock) to a 3 mM ATP final concentration in an Eppendorf tube. The reaction mixture consisted of 5 μg/ml LMCA1, 1.1 mM CaCl_2_, 0.1 mM EGTA, 50 mM Tris, 200 mM KCl, 10 mM MgCl_2_, 20% v/v glycerol, 1 mM BME, 0.25 mg/mL C_12_E_8_, pH=7.6 and the sample reconstituted in nanodiscs was in a reaction mixture without detergent and glycerol (5 μg/ml LMCA1, 1.1 mM CaCl_2_, 0.1 mM EGTA, 50 mM Tris, 200 mM KCl, 10 mM MgCl_2_, 1 mM BME, pH=7.6). The negative control contained 1.1 mM EGTA and no CaCl_2_. After 2-8 min 50 μl of the reaction mix was terminated in a 96-well plate with 50 μl freshly prepared stop solution (140 mM ascorbic acid, 5 mM (NH_4_)_2_MoO_4_, 0.1% w/v SDS, 0.4 M HCl) in each well and incubated for 15 min. The blue color formed was stabilized with 75 μl arsenite solution (150 mM sodium arsenate, 70 mM sodium citrate, 350 mM acetic acid). After 30 min incubation, the absorbance was measured at 860 nm and the activity was calculated.

### Cryo-EM grid preparation and data collection

The sample mix (0.7 mg/ml saposin-solubilized G_4_-LMCA1, 50 mM Tris, 150 mM KCl, 1 mM CaCl_2_, 10 mM MgCl_2_, 0.0015% LMNG, pH=7.6) was supplemented with 1 mM ATP and incubated at room temperature for 2-10 minutes. 3 µl were added to a freshly glow-discharged grid (45 s at 15 mA), which was subsequently blotted at 20 °C and 90% humidity for 4 seconds. CF-2/2-3Cu (Protochips) were used, except for CF-1.2/1.3-3Cu in one case. Blotting and subsequent plunge freezing into liquid ethane were carried out on an EM GP2 plunge freezer (Leica) with one-two filter papers. Data were collected on a Titan Krios G3i microscope (EMBION Danish National cryo-EM Facility – Aarhus node) operated at 300 KeV equipped with a BioQuantum energy filter (energy slit width 20 eV) and K3 camera (Gatan). Movies were collected using aberration-free image shift data collection (AFIS). A nominal magnification of 130,000x was used, resulting in a pixel size of 0.647 Å^2^/px with a total dose of 59.0 e^-^/Å^2^ (grid 1+2) and 58.5 e^-^/Å^2^ (grid 3+4). The movies were fractionated into 53 frames (1.13 e^-^/Å^2^ per frame) at a dose rate of ∼18 e^-^/px/s and a 1.4 s exposure time per movie. A nominal defocus range of -0.8 to -2.0 µm was used.

### Cryo-EM image processing

Figure S2 shows the processing pipeline for image processing. Movies were motion corrected^36^ and CTF estimated^37^ using CryoSPARC (v3 and v4)^38^. Particles from grid 1 were picked using a circular blob and aligned by 2D classification. Small subsets of the 2D classes were selected and used to generate *ab initio* volumes. One protein-like and one-two junk volumes were used to select protein particles from junk in multiple rounds of heterogenous refinement. This particle stack was used as template for template picking in all micrographs, which were merged from four grids to extract 3,001,118 particles. Again, particles from selected 2D classes were used to generate *ab initio* volumes for separating protein from junk particles in multiple rounds of heterogeneous refinement. Global CTF refinement followed by non-uniform refinement^39^ of 101,034 particles generated a volume that was used to manually generate a mask in ChimeraX^40^. This mask was used in 3D variability ^41^ in cluster mode. One of the clusters provided a volume that was finally used for non-uniform refinement.

### Homology modeling

A homology model for the [Ca]E1P-ADP state of LMCA1 was made using an online version of Modeller (www.salilab.org/modeller)^42^. A sequence alignment of LMCA1 and SERCA performed in MUSCLE ^43^ was used as an input file together with the structure of SERCA stabilized with Ca^2+^, ADP and AlF_x_ (PDB: 15t5)^12^.

### Model building, refinement, and validation

A fusion model was created by using the cytosolic domain from the crystal structure of G_4_-LMCA1 E2P (pdb: 6zhh)^25^ and the TM domain of the LMCA1 homology model for the [Ca_2_]E1P-ADP state. The model was manually docked in the cryo-EM volume and fitted to the map with geometry restraints using Namdinator^44^.

Real space refinement of the structure was done in Phenix ^45^, and model building and analysis were performed in Coot^46^ (Table S2). A DeepEMhancer^47^ map further guided model building.

## Acknowledgements

We dedicate this paper to the memory of the late professor Jesper Vuust Møller (1938-2023), who founded studies of Ca2+-ATPase at Aarhus University and with whom we have had numerous fruitful discussions and collaborations on Ca^2+^-ATPases. We are grateful to Jens Peter Andersen for critical reading of the manuscript. The authors would like to thank Anna Marie Nielsen for technical assistance, to Josephine Karlsen Dannersø and the staff of the EMBION cryo-EM facility for help with sample preparations, optimisation and data collection, and to Samuel Hjort-Jensen and Jesper Karlsen for help with data processing and structure determination using the EMCC facilities at Aarhus University for scientific computing. This work was supported with a PhD scholarship from the Boehringer Ingelheim Fonds to S.B.H., a professorship grant from the Lundbeck Foundation (R310-2018-3713) to P.N. and a project 2 grant from the Danish Fund for Independent Research (7014-00328B) to M.K. and P.N. Research infrastructure was supported with a grant from the Danish Ministry for Research and Higher Education (EMBION cryo-EM facility, grant no. 5072-00025B), and the Novo Nordisk Foundation (ICE-T facility, grant no. NNF20OC0060483)

