## Supplemental Information for "Structure of a Ca^2+^ bound phosphoenzyme intermediate in the inward-to-outward transition of Ca^2+^-ATPase 1 from *Listeria monocytogenes*"

Supplementary Information

**Table S1 pairwise comparison of C_α_’s of G_4_-LMCA1 [Ca]E2P to LMCA1 and SERCA structures.**

| **Molecule 1** | **Molecule 2** | **Full molecule** | ***Cytosolic domains** | ****TM domain** |
| --- | --- | --- | --- | --- |
| G_4_-LMCA1  [Ca]E2P | SERCA [Ca]E1- AlF_x_-ADP (pdb: 1t5t) | 7.6 Å | 12.2 Å | 2.1 Å |
|  | LMCA1 E2-BeF_x_ (pdb:6zhf) | 4.2 Å | 2.9 Å | 3.3 Å |
|  | SERCA E2-BeF_x_ (pdb:3b9b) | 3.5 Å | 1.9 Å | 5.0 Å |
|  | LMCA1 E2-AlF_x_ (pdb:6zhg) | 4.7 Å | 3.0 Å | 3.4 Å |
|  | SERCA E2-AlF_x_ (pdb:39br) | 5.0 Å | 3.1 Å | 4.2 Å |

* The cytosolic domains are defined as:

LMCA1: Residue 1-35 and 112-225 and 314-655
SERCA: Residue 1-37 and 124-241 and 331-735

** The TM domains are defined as:

LMCA1: Residue 36-111 and 226-313 and 656-880

SERCA: Residue 38-123 and 242-330 and 736-994

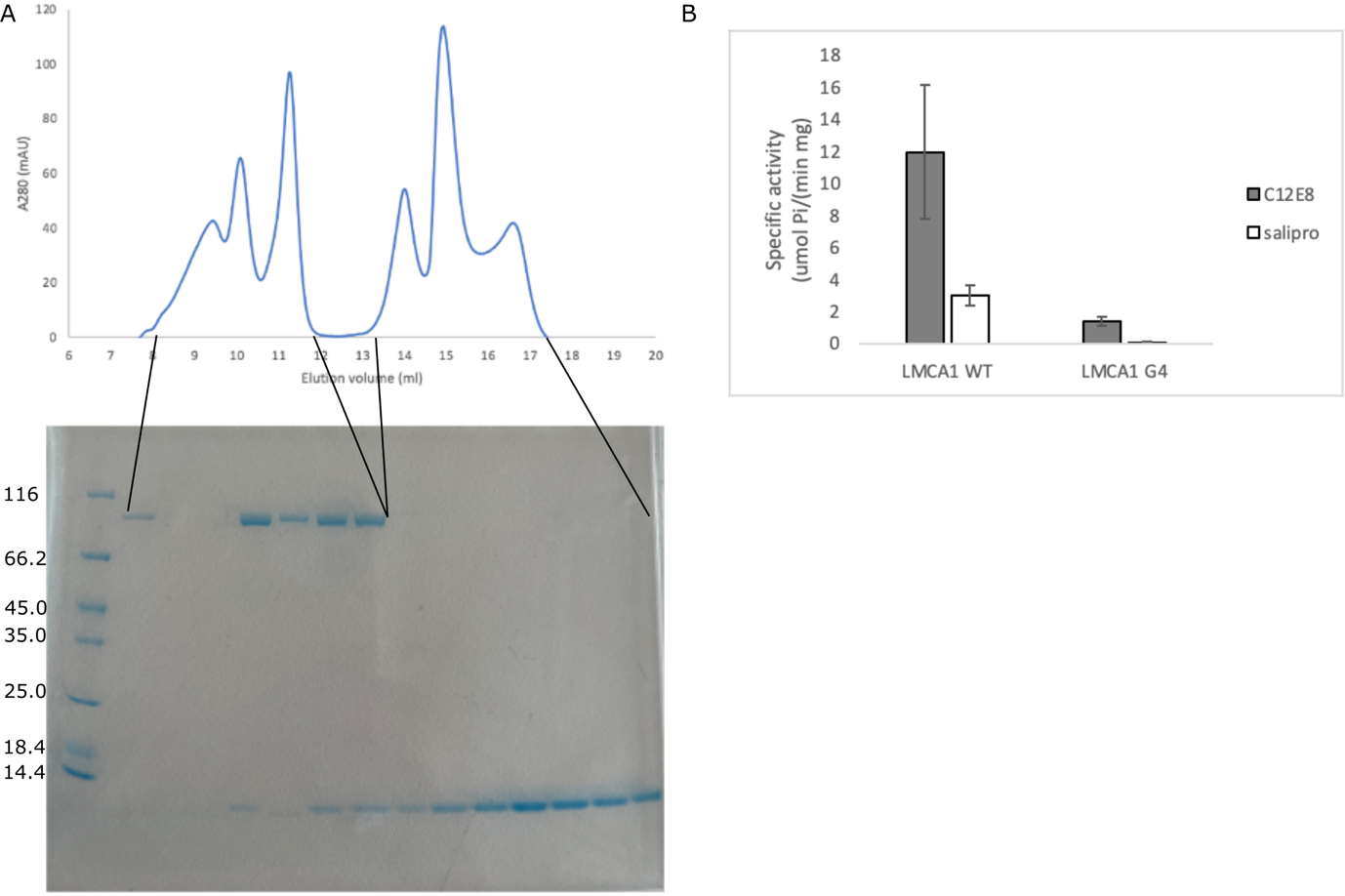

**Figure S1: Salipro reconstitution and activity.** (**A**) Chromatogram of salipro reconstitution of LMCA1. The peak eluting at 11-12 ml contains monomeric LMCA1 in nanodiscs. (**B**) Specific ATPase activity is obtained by linear regression of a time-course measurement.

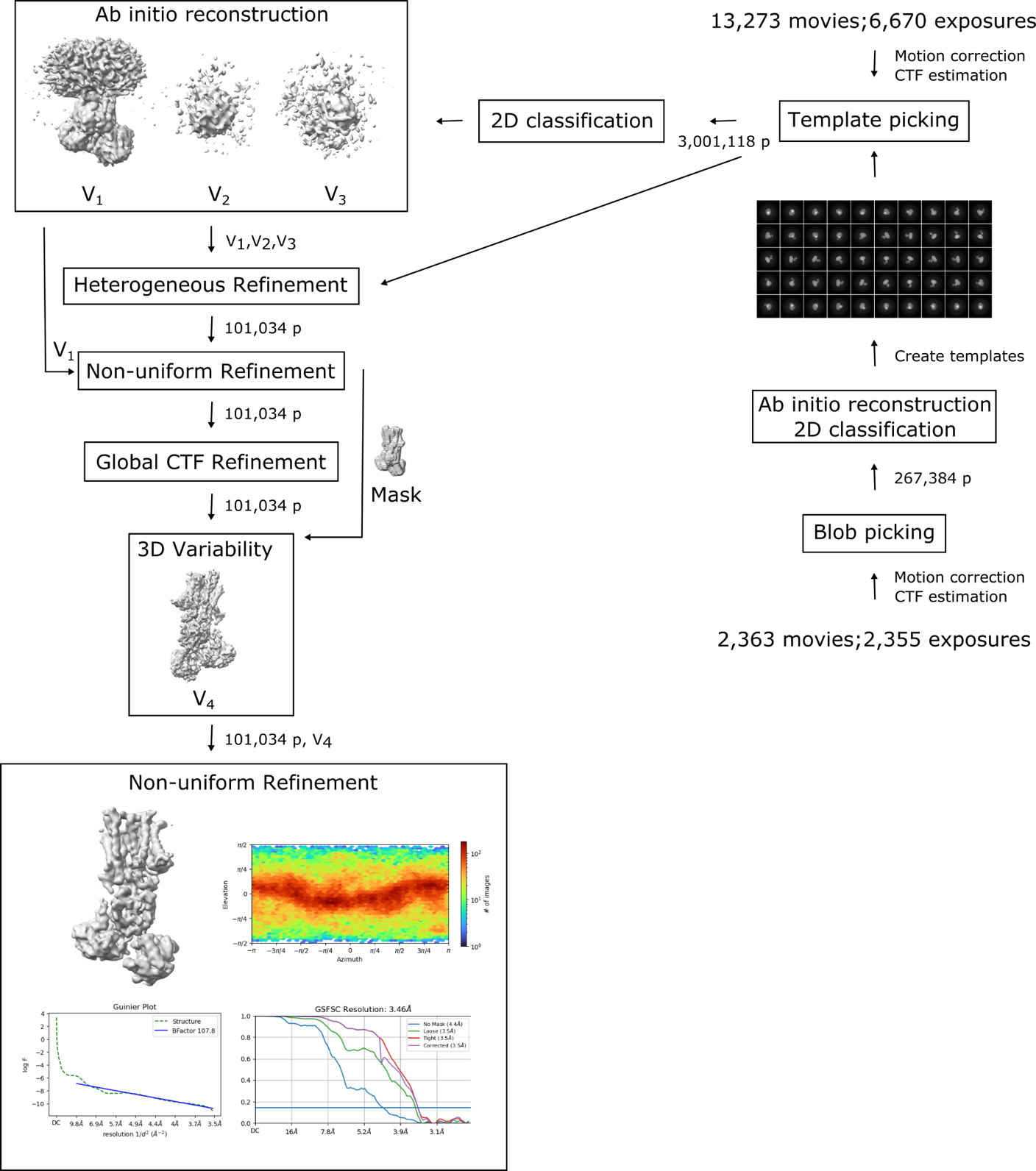

**Figure S2: Data processing pipline for cryo-EM single particle analysis.**

**Table S2. Data collection and refinement for cryo-EM single particle analysis.**

| **Data collection and processing** |  |
| --- | --- |
| Magnification | 130,000x |
| Voltage (kV) | 300 |
| Microscope | Titan Krios G3i |
| Camera | Gatan K3 |
| Physical pixel size (Å/pix) | 0.647 |
| Electron exposure (e-/Å^2^) | 59.0 (grid 1+2), 58.5 (grid 3+4) |
| Defocus range (µm) | -0.8 to -2.0 |
| Total number of movies | 13,273 |
| Initial particles | 3,001,118 |
| Final particles | 101,034 |
| Symmetry imposed | C1 |
| Map resolution (Å) | 3.46 |
| FSC threshold | 0.143 |
| **Refinement** |  |
| Initial model used | Fusion between ‘pdb: 6zhh’ and LMCA1 homology of ‘pdb: 1t5t’ |
| Atoms | 6720 |
| Residues | 880 |
| Ligands | 3 (Ca2+, Mg2+, -PO_3_^-^) |
| Average B factor (Å^2^) |  |
| Bond length RMSD (Å) | 0.003 |
| Bond angle RMSD (°) | 0.699 |
| MolProbity score | 2.07 |
| Clashscore | 14.46 |
| Ramachandran outliers (%) | 0 |
| Ramachandran allowed (%) | 5.94 |
| Ramachandran favored (%) | 94.06 |
| PDB code | xxxx |

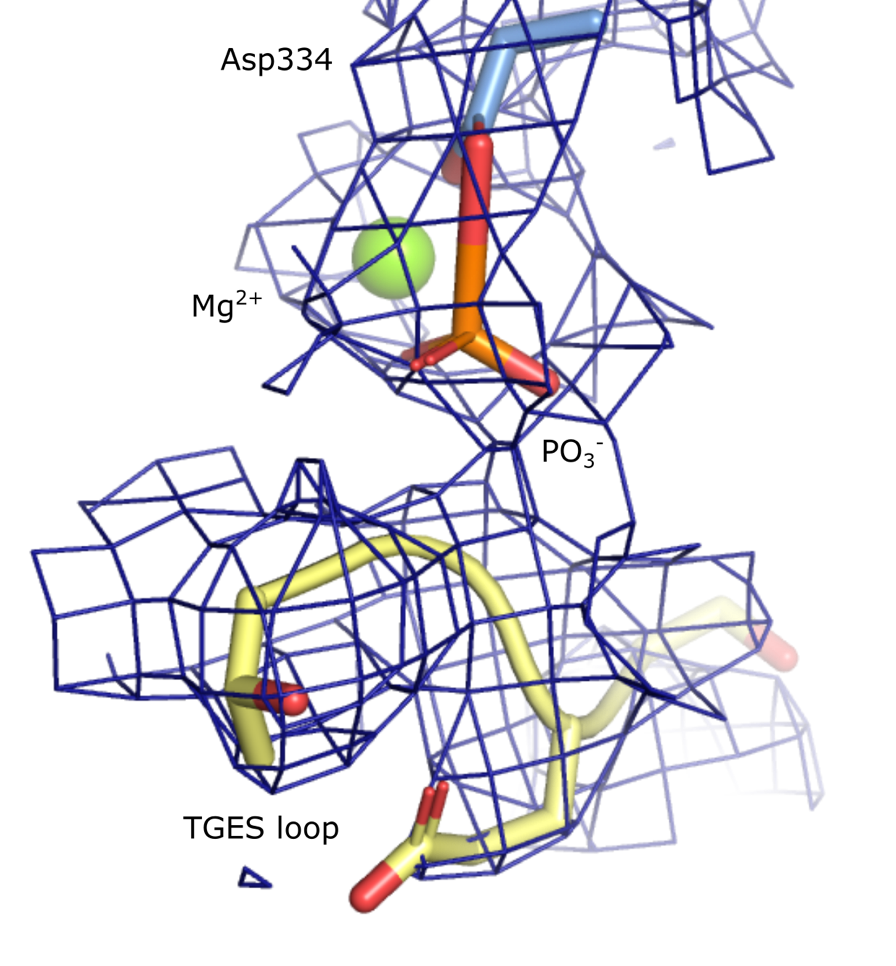

**Figure S3: Density for phosphorylation.** The catalytic aspartate (Asp334) is phosphorylated, and the phosphate group is coordinated by a Mg^2+^ ion. The TGES loop is protecting the phosphorylation site. Density is only shown for the TGES loop in yellow, Asp334 in blue, the phosphorylation and the Mg^2+^ ion.

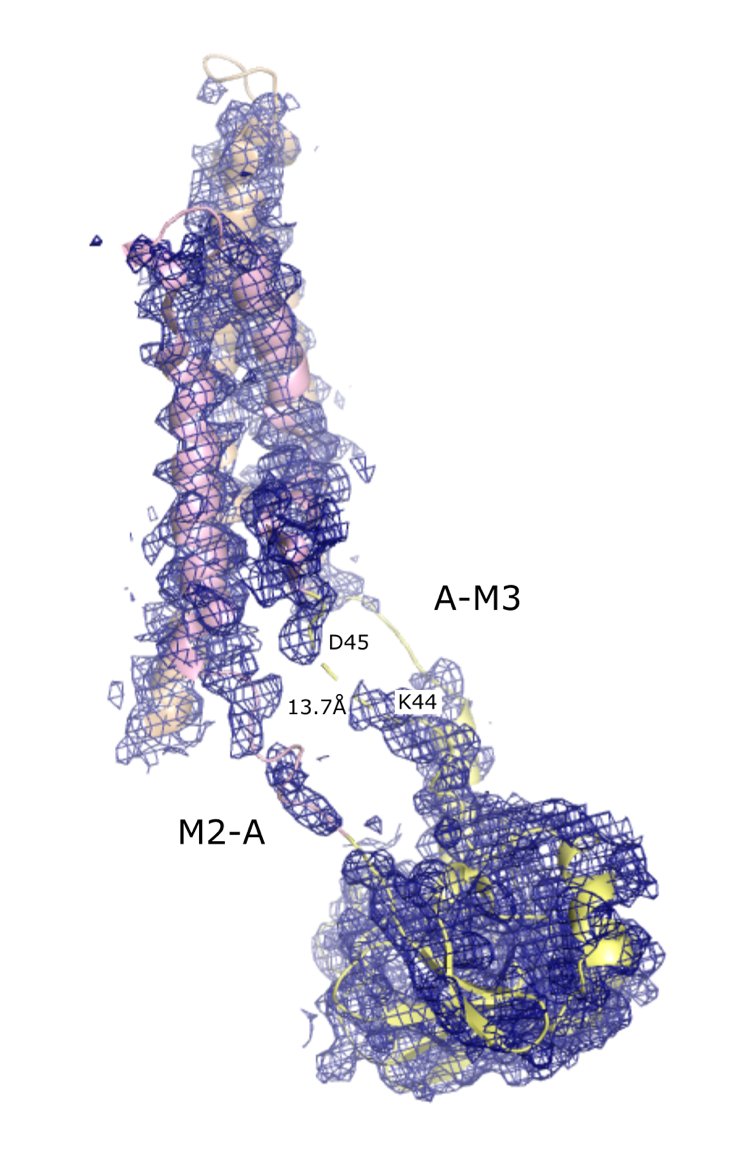

**Figure S4: Density for A domain linkers in [Ca]E2P.** The density is shown for M1-4 and the A domain of the [Ca]E2P. The model is represented as cartoon and colored according to Figure 1. The break in linker A-M1 is indicated with numbered residues and the C_α_ Lys44 - C_α_Asp45 is measured.
